# Visual Search and ASD: Basic vs. Superordinate Category Search

**DOI:** 10.64898/2026.11.17.767228

**Authors:** Safa’a Abassi Abu Rukab, Cory Shulman, Shaul Hochstein

## Abstract

We study visual search and categorization among individuals with Autism Spectrum Disorder (ASD) and a neuro-typical comparison group (NT). Individuals with ASD often exhibit distinct sensory processing patterns and may face challenges in efficiently detecting and categorizing visual stimuli. They may also process categories differently compared to NT groups, reflecting variations in typical sensory processing mechanisms. Based on our previous findings, this study employs visual search categorization tasks, testing reaction time and set-size slopes for different target categories. Participants included 26 individuals with ASD and 11 NT individuals, with similar mean ages and mean IQ Block Design subtest scores. They performed basic-level and superordinate-level visual search, as well as a categorization task. In the visual search task, participants identified targets within arrays of varying set sizes (4, 16, 36, 64). Face detection remained rapid across groups, and Basic-level search was similar in ASD and NT groups (mean set-size slopes: 48.7 and 45 ms/item, respectively). In contrast, superordinate search was much slower in ASD (99 ms/item) compared to NT (70.8 ms/item), reflecting increased cognitive demands. Object typicality influenced performance, with less typical items being slower to both categorize and detect. These findings suggest that basic-level processing is relatively preserved in ASD, whereas superordinate categorization imposes greater demands. The results highlight the role of typicality and support differences in perceptual and cognitive processing in ASD. Our findings contribute to better understanding visual cognition in autism and may inform strategies for improving conceptual generalization and visual learning.

## Introduction

Autism spectrum disorder (ASD) is a neurodevelopmental condition defined by persistent difficulties in social communication and interaction, along with restricted or repetitive behaviors, interests, or activities. These characteristics emerge early in development and affect daily functioning. Individuals with autism may also respond differently to sensory experiences, showing increased or reduced sensitivity to sensory input or a strong interest in particular sensory features of their surroundings (American Psychiatric Association, 2013). Individuals with autism exhibit distinct sensory processing patterns compared to non-autistic individuals, impacting auditory, visual, and cognitive functions (Dellapiazza et al., 2018; DuBois et al., 2017; Burns et al., 2017; Bölte et al., 2019; Shulman et al., 2020a,b).

Understanding how people with ASD attend to their surroundings requires examining how relevant visual information is selected among competing objects. One way to investigate this process is through visual search, in which a person locates a target among distractors. The ease of finding that target depends partly on the visual characteristics that distinguish it from the surrounding items (Treisman & Gelade, 1980; Wolfe, 2021). Enhanced visual search has been reported in participants with autism. For example, Plaisted et al. (1998) found that children with autism located conjunction targets faster than comparison children. However, findings vary across tasks and samples, suggesting that visual-search performance in autism depends on both what participants are searching for and how the task is structured (Doherty et al., 2018; Joseph et al., 2009).

In their feature-integration theory, Treisman and Gelade (1980) distinguished between feature search and conjunction search. In feature search, also called odd-one-out or pop-out search, a target may stand out because of a single distinctive property, such as its color, allowing it to be detected rapidly with little effect of the number of items in the display. In conjunction search, identifying the target requires combining properties, such as color and shape, and such search typically becomes more time-consuming as the number of items increases. This number, referred to as set size, allows researchers to examine how search performance changes with increasing display demands. The distinction between feature and conjunction search is also evident in children and it develops with age. For example, Donnelly et al. (2007) compared children aged 6–7 and 9–10 years with young adults and found that children were slower in both odd-one-out and conjunction searches, with greater effects of set size in the youngest group. These findings suggest that differences in visual search performance should be interpreted in relation to both participants’ age and the demands of the task.

Reverse hierarchy theory offers a framework for understanding rapid category detection (Hochstein & Ahissar, 2002). It proposes that the whole is perceived first due to explicit perception initially accessing only high-level representations. Later explicit perception returns to lower areas via the feedback connections, to provide and integrate into conscious vision with scrutiny, detailed information about the scene. Hochstein (2020) related this distinction to global and focused attention: global attention supports an overall impression of a scene, whereas focused attention allows closer examination of its details. Reverse hierarchy theory predicts that pop-out search is high level and should include face-search. Indeed, Hershler and Hochstein (2005) found a face-search advantage among heterogeneous photographs of objects, and later work showed that faces were still detected fairly well at larger eccentricities and with the presence of flankers (Hershler et al., 2010). The detection of a category is therefore distinct from the identification of one of its members.

This distinction between detection and identification was also examined in our previous study of visual search in participants with ASD and non-autistic participants (Abassi Abu Rukab et al., 2022). The study included 23 participants with ASD and 23 non-autistic participants matched for age and IQ Block design subtest scores, who searched for human faces and other target categories among arrays of 4–64 photographs. Although ASD participants were generally slower, both groups showed a similar pattern across target categories: human faces were found fastest and with the shallowest search slopes. Face-search slopes were significantly steeper in the ASD group (18 vs. 11 ms/item), but search remained highly efficient in both groups, approaching pop-out performance. In contrast, searching for the broader animal-face category was less efficient, particularly for ASD participants. These findings suggest that rapid face detection can be relatively preserved despite reported difficulties in face recognition, consistent with a distinction between vision at a glance and vision with scrutiny. They also raise the question of how broadly-defined target categories influence search efficiency in autism.

How the target is named also matters. A basic-level label, such as “cat” or “car,” refers to a relatively specific basic category, whereas a superordinate label, such as “animal” or “vehicle,” includes a broader range of objects (Rosch et al., 1976; Taniguchi et al., 2020). This distinction concerns category level and is separate from the distinction between feature and conjunction search. Schmidt and Zelinsky (2009) showed that more specific target cues provide more effective guidance during visual search. Maxfield and Zelinsky (2012) showed that category level can affect both how attention is directed during search and how quickly an object is confirmed as the target. These findings suggest that the way a target is named can influence both finding a likely object and deciding whether it matches the target description.

Evidence from young children suggests that categorical search is particularly relevant to understanding visual performance in autism. Doherty et al. (2018) examined search in three-year-old children, including children with an increased familial likelihood of autism. Higher autism symptom severity was associated with poorer categorical search, rather than an overall search advantage. Although their study examined symptom differences in a younger sample, it highlights the importance of considering whether participants are searching for a specific kind of object or for members of a broader category.

Taniguchi et al. (2020) found that the visual properties associated with categorization differed between basic and superordinate levels, and that categorization depended partly on whether the category name appeared before or after the image. Similarly, Tiedemann et al. (2022) showed that the structure and arrangement of object parts affected whether unfamiliar shapes appeared more like animals or plants. Together, these findings suggest that visual information also contributes to superordinate judgments, so differences in performance between category levels may reflect both perceptual and semantic demands.

These findings raise a broader question: how does search efficiency change when the target is defined by a category that includes visually different objects? Comparing basic and superordinate search across several everyday categories allows us to examine whether the additional cost of a broader target definition differs between participants with or without autism.

### The present Study

The present study examined how category level affects visual-search efficiency in participants with and without autism, and whether performance in visual search is related to performance in categorization.

In the visual-search task, we examined visual search in 26 participants with autism spectrum disorder (ASD) and 11 neurotypical (NT) participants. Participants searched for target images presented among arrays of 4, 16, 36, or 64 items on a touchscreen. Search was tested at both the basic level, such as car, cat, or apple, and the superordinate level, such as vehicle, animal, or food. We also included a face-search condition, in which participants searched for a human face among non-face distractors. Reaction times across the different set sizes were used to examine search efficiency and calculate search slopes. Figure (1) shows examples of the stimuli used (musical instrument, apple, cat, food).

**Figure 1.**
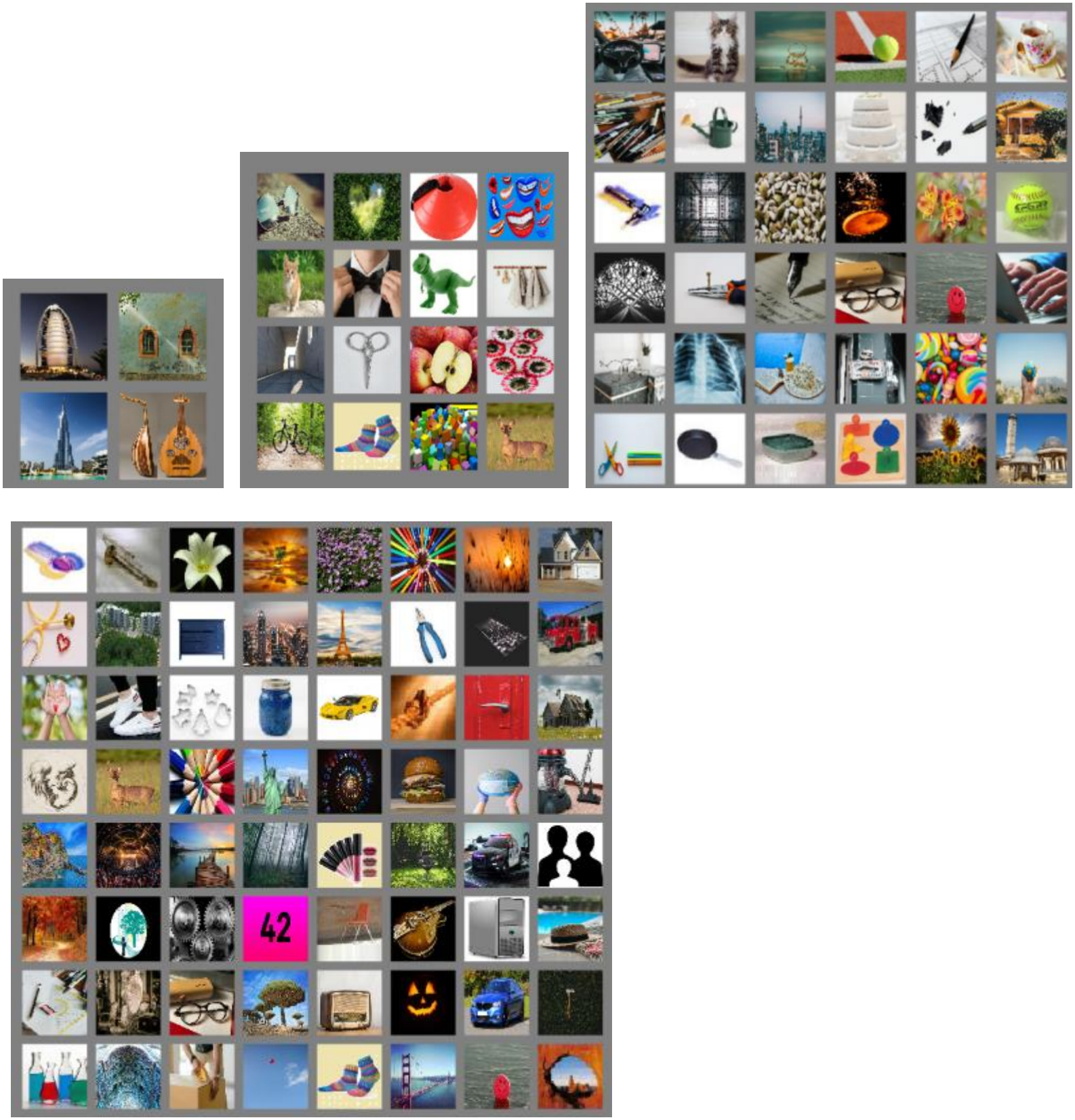
Example target categories and set sizes: (A) musical instrument, 4 items; (B) apple, 16 items; (C) cat, 36 items; and (D) food, 64 items. Each display contains one target among distractors.

In a separate categorization task, participants heard a category name in Arabic and then saw two images. They were asked to choose the image that represents the named category. Reaction times for the same images were compared across the two tasks to examine whether images selected more quickly in the categorization task tended to be also found more quickly during visual search.

## Methods

### Participants

A total of 37 participants took part in the study, including 26 participants with autism spectrum disorder (ASD) and 11 neurotypical (NT) participants. The ASD group consisted of 22 males and 4 females, while the NT group included 8 males and 3 females.

The mean (±SE) age was 16.64±0.49 years in the ASD group and 15.06 ±0.66 years in the NT group. An independent-samples t-test indicated that the age difference did not reach statistical significance (p = .069).

Visuospatial performance was assessed with the Block Design subtest from either the WISC-IV (Wechsler, 2003; children below age 17) or the WAIS-III (Wechsler, 1997; from age 17). Raw scores included time-bonus points. The ASD group obtained a mean score of 32.17± 2.85, compared with 34.18± 3.07 in the NT group. The reported comparison of these raw scores was not statistically significant (p = .635). These scores describe performance on the administered subtests and should not be interpreted as full-scale IQ or evidence that the groups were matched in general cognitive ability.

Autism-related traits were assessed using the 50-item Autism Spectrum Quotient (AQ; Baron-Cohen et al., 2001), obtained from the Psychology Tools website (Psychology Tools, n.d.). The questionnaire was completed under the supervision of the experimenter based on information provided by school staff and caregivers. AQ scores were significantly higher in the ASD group (M = 27.65±1.23) than in the NT group (M = 11.45±0.99; p < .001).

Participants in the ASD group were recruited from a special education school in East Jerusalem that serves students formally diagnosed with autism spectrum disorder according to national special education committee criteria. The NT group was recruited from nearby mainstream community schools. Participants with motor impairments, as identified by the school occupational therapist, were excluded from the study.

### Ethics and approval

All procedures were approved by the Chief Scientist of the Israel Ministry of Education and authorized by the school administration. Written informed consent was obtained from parents or legal guardians of all participants.

### Stimuli

The stimuli consisted of photographs of everyday objects representing basic-level and superordinate-level categories. Basic-level categories included objects such as cat, car, apple, spoon, and keyboard, with corresponding superordinate categories including animal, vehicle, food, kitchen tool, and musical instrument. A human-face search condition was also included, in which photographs of real human faces served as targets among non-face distractor images. The same stimulus sets were used for all participants.

For the visual-search task, one target image was presented among distractor images in arrays containing 4 (2 × 2), 16 (4 × 4), 36 (6 × 6), or 64 (8 × 8) items. For the categorization task, two images were presented simultaneously; in both cases search followed an auditory category cue in Arabic.

### Procedure

Participants were tested individually in a quiet room at their school (ASD) or familiar environment (NT). All tasks were administered by a trained experimenter following a standardized protocol, and instructions were given verbally with comprehension checks prior to task initiation.

In the first testing session, participants began with the face-search task, in which they located and touched a human face among non-face distractors. They then completed one basic-level search task (e.g., “car”), followed immediately by the corresponding superordinate-level task (e.g., “vehicle”). Subsequent sessions followed the same basic-to-superordinate sequence using a different category pair each time. The face-search task was administered only once, at the beginning of the first session. Sessions took place on separate days, approximately one to two days apart.

Each category included 40 trials, with 10 trials at each set size (4, 16, 36, and 64 items). Each trial began with a fixation cross (1 s exposure), followed by stimulus presentation. Participants were instructed to respond as accurately and quickly as possible. Feedback was provided via auditory signals: a success sound (high-pitch tone) for correct responses and an error sound (low-pitch tone) for incorrect responses. Participants were allowed up to two attempts per trial, and trials had a maximum duration of 30 seconds before automatic skipping. Data analyses excluded two-attempt trials.

For the categorization task, participants completed a two-alternative forced-choice (2AFC) word-picture matching paradigm. Each trial began with a spoken category cue (in Arabic), followed by presentation of two images side by side: a target belonging to the named category and a distractor, an image of an object from a different category. Participants were instructed to select the image that best represented the named category as quickly and accurately as possible. The task consisted of two blocks of 100 trials each, yielding 200 trials in total. Five categories were included: animals, food, vehicles, kitchen tools, and musical instruments, with 40 trials per category. Each target image appeared once, paired with a distractor, and trials from the different categories were intermixed.

Response time and accuracy were recorded for each trial.

The structure of the total number of sessions and the orders of the tasks per participants remained consistent across participants and groups.

### Apparatus

Displays were presented to the participants on a HP laptop with a 14-in. touchscreen (1,920 × 1,080 pixels). The laptop screen could be rotated vertically around so that the keyboard was invisible. All participants used the same touchscreen laptop computer for these experiments, so that conditions were equivalent for all.

Each stimulus image measured approximately 1.80 × 1.80 cm on the screen, images were arranged in regular grids with approximately 2.5 mm spacing between adjacent items. Stimulus size and spacing were kept constant across trials and experimental conditions. The laptop rested on the participant’s lap or a desk, approximately 40–50 cm from the participant’s face.

### Data analysis

Data were organized in Microsoft Excel to calculate search slopes, slope differences, and normalized reaction times. Accuracy was calculated as the percentage of trials answered correctly within each group and search condition, using the corresponding number of completed trials.

Reaction times from incorrect trials, including trials requiring a second attempt, were excluded from reaction-time analyses.

Participant-level basic and superordinate search slopes were analyzed using a mixed ANOVA with search level (basic or superordinate) as the within-participant factor and group (ASD or NT) as the between-participant factor. Superordinate search cost was calculated by subtracting each participant’s basic-level slope from their superordinate-level slope. Face-search results were described separately. Group means were plotted with 95% confidence intervals calculated from the standard error and the *t* distribution.

Pearson correlations between basic- and superordinate-level slopes were tested separately within each group using one-tailed tests, based on the positive relationship predicted in advance. Pearson’s r and R^2^ were reported to describe associations. Differences between the groups’ correlation coefficients were not tested. Figures were prepared using Microsoft Excel, IBM SPSS Statistics, and Python. Linear regressions summarized the relationships shown in the scatterplots.

## Results

### Visual search across set sizes

Overall accuracy was high in both groups (ASD: 96.3%; NT: 98.2%). Basic-level accuracy was descriptively similar across groups (ASD: 99.2%; NT: 99.1%), whereas superordinate-level accuracy was lower in the ASD group (92.8%) than in the NT group (96.9%).

Reaction times increased with set size in both groups across basic-level, superordinate-level, and face search (Figure 2). The set-size increase was greatest for superordinate-level search and smallest for face search, with basic-level search falling between the two. In the NT group, the slopes were

**Figure 2.**
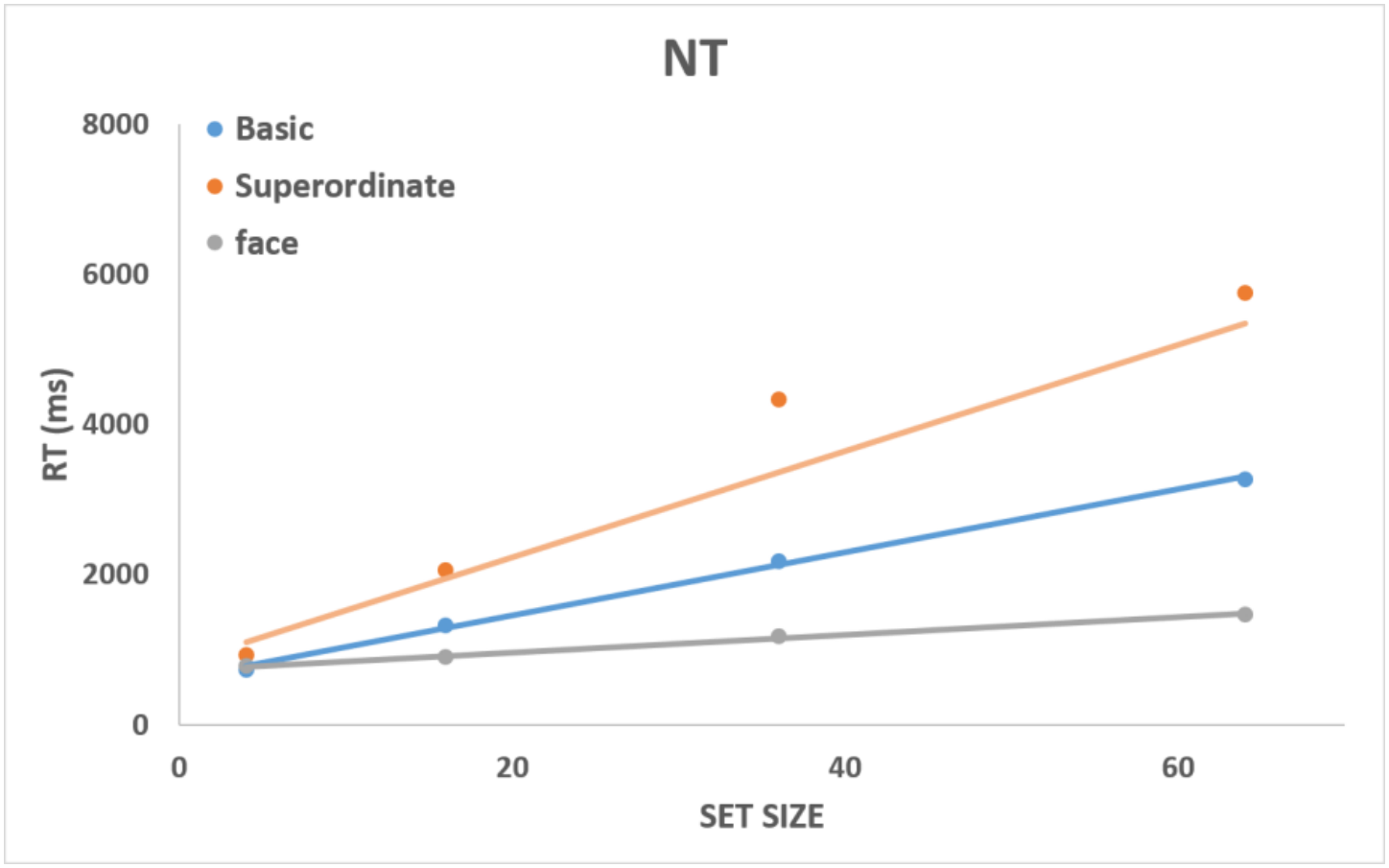
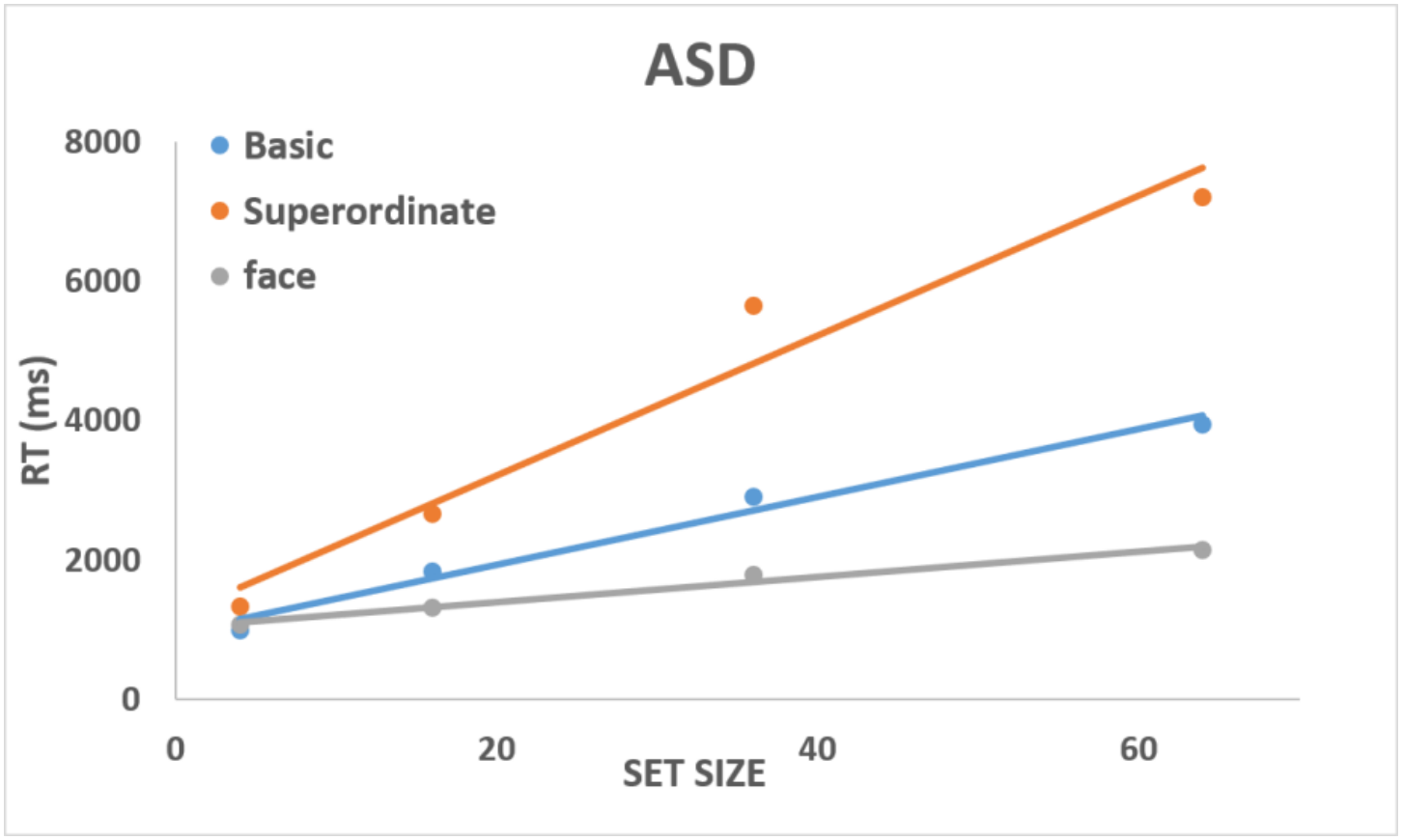
Mean reaction time (milliseconds) as a function of set size in the NT group (top) and ASD group (bottom). The three series represent basic-level, superordinate-level, and face search.

45.0 ms/item for basic-level search, 70.8 ms/item for superordinate-level search, and 11.7 ms/item for face search. In the ASD group, the corresponding slopes were 48.7, 99.0, and 18.0 ms/item. Thus, slopes were consistently greater for the ASD group, but both groups showed the same ordering: reaction times increased most sharply during superordinate-level search and least sharply during face search.

### Basic and superordinate search slopes

To examine differences in search efficiency, individual participants’ slopes were entered into a mixed ANOVA with search level (basic or superordinate) as the within-participant factor and group (ASD or NT) as the between-participant factor. (Figure 3). The main effect of search level was significant, F(1, 35) = 89.08, p < .001, ηp^2^ = .718, with steeper slopes for superordinate-level search. There was also a significant main effect of group, F(1, 35) = 4.92, p = .033, ηp^2^ = .123, indicating steeper slopes overall in the ASD group. The Search Level × Group interaction was significant, F(1, 35) = 9.25, p = .004, ηp^2^ = .209. The increase from basic to superordinate search was greater in the ASD group (50.30 ms/item) than in the NT group (25.78 ms/item) and the difference between groups was greater for superordinate level search.

**Figure 3.**
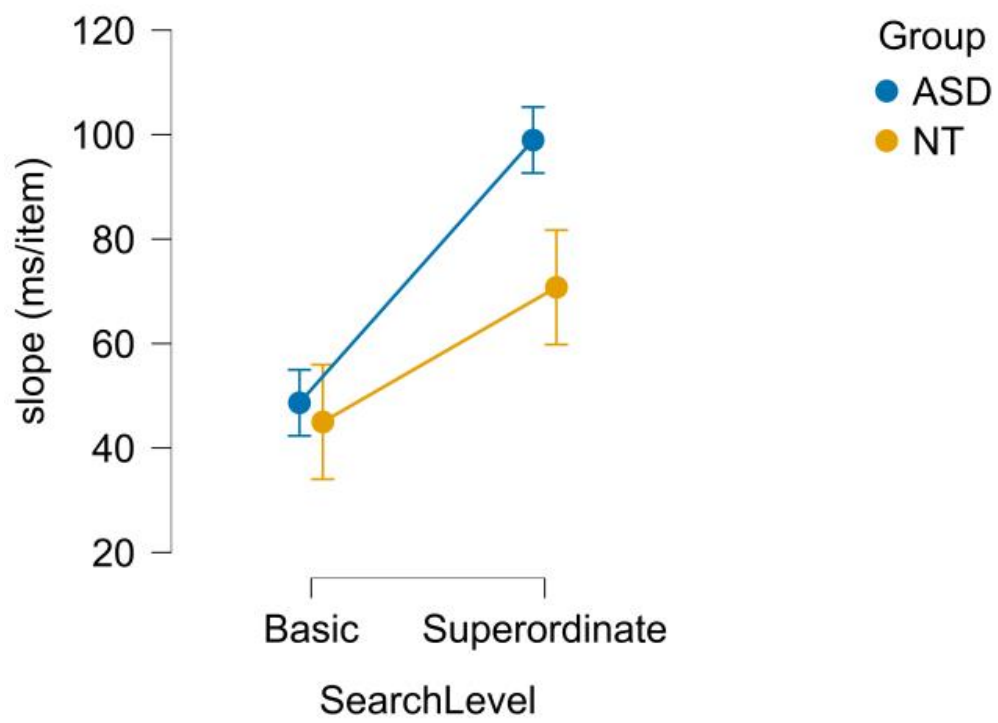
Group mean slopes, with 95% confidence intervals

### Individual differences in search slopes

The paired slopes show how performance changed from the basic to the superordinate level for each participant (Figure 4). Superordinate-level slopes were generally steeper, resulting in positive difference scores for most participants. This increase was larger on average in the ASD group, although its magnitude varied considerably within both groups.

**Figure 4.**
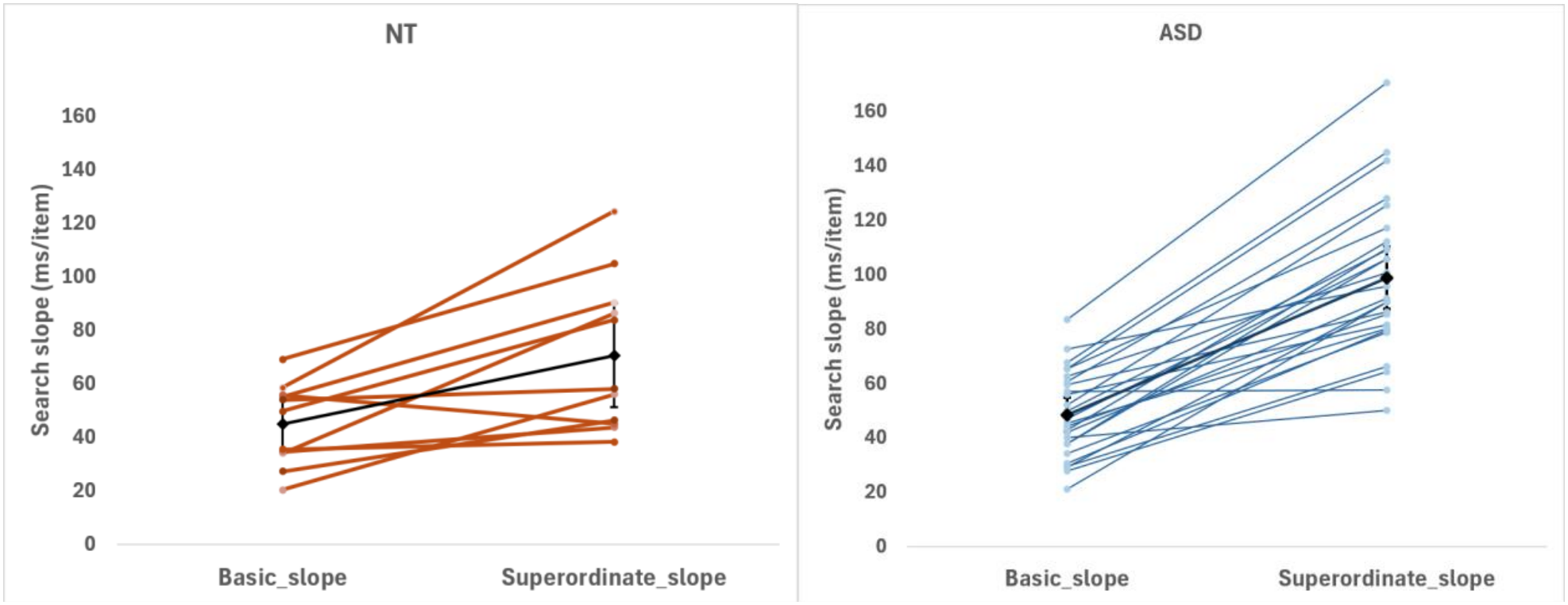
Individual basic- and superordinate-level search slopes in the NT group (top) and ASD group (bottom). Each colored line connects the two slopes for one participant. Black diamonds indicate group means.

### The increase from basic to superordinate search

The individual difference scores, subtracting the basic search slope from the superordinate search slope for each participant, further illustrate the larger increase in the ASD group (Figure 5). Most scores were positive, indicating that adding items to the display had a greater effect on reaction time during superordinate-level search. The distributions show substantial variation among participants in both groups.

**Figure 5.**
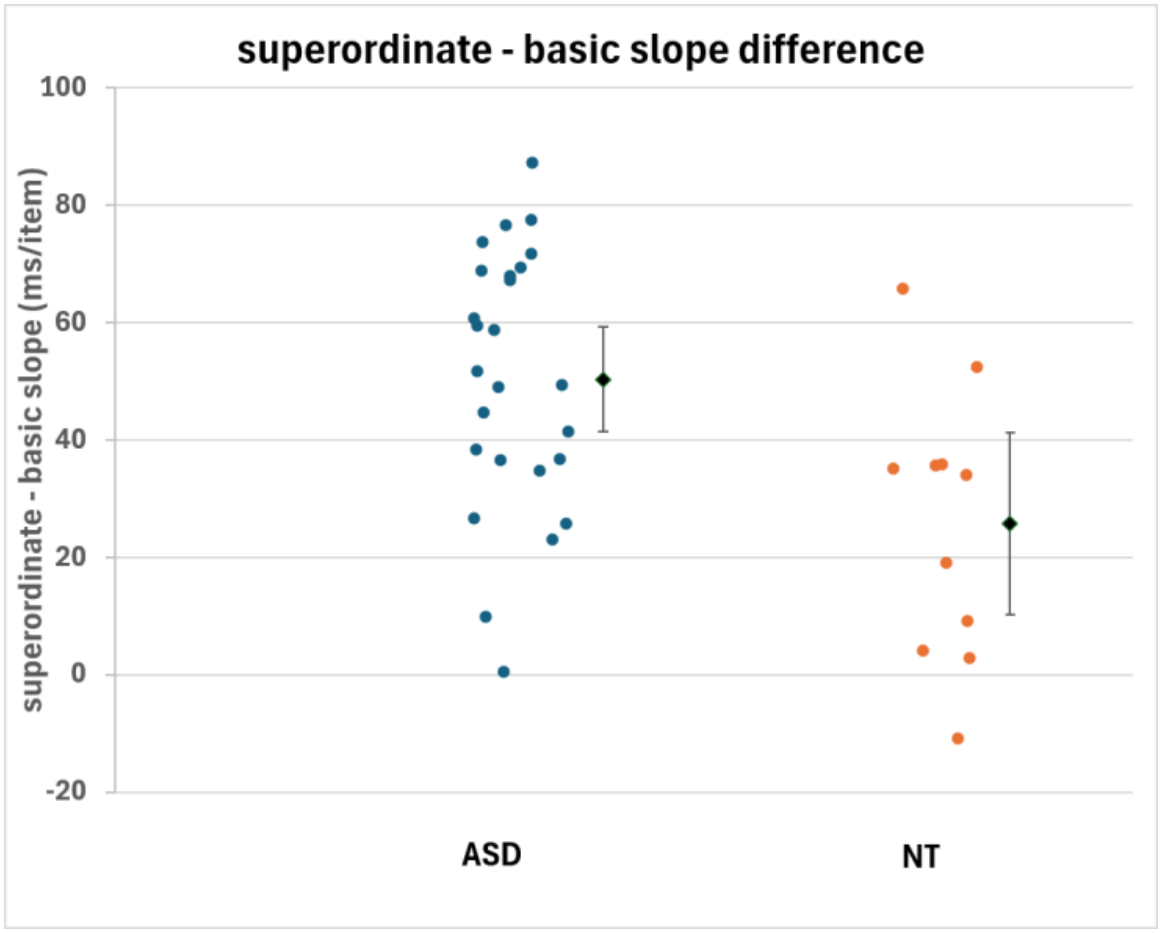
Differences between superordinate- and basic-level visual-search slopes in the ASD and NT groups. Dots represent individual participant’s superordinate minus basic slope. Black diamonds indicate group means, with error bars showing 95% confidence intervals calculated from the standard error using the *t* distribution.

### Correlations between basic and superordinate slopes

Participants with steeper basic-level search slopes also tended to have steeper superordinate-level slopes (Figure 6). Positive correlations of similar magnitude were found in the ASD group, *r*(24) = .624, *p* < .001, and the NT group, *r*(9) = .600, *p* = .025. Both correlations were statistically significant using one-tailed tests, consistent with the prespecified prediction of a positive relationship. Thus, participants whose search times increased more sharply with set size during basic-level search tended to show the same pattern during superordinate-level search.

**Figure 6.**
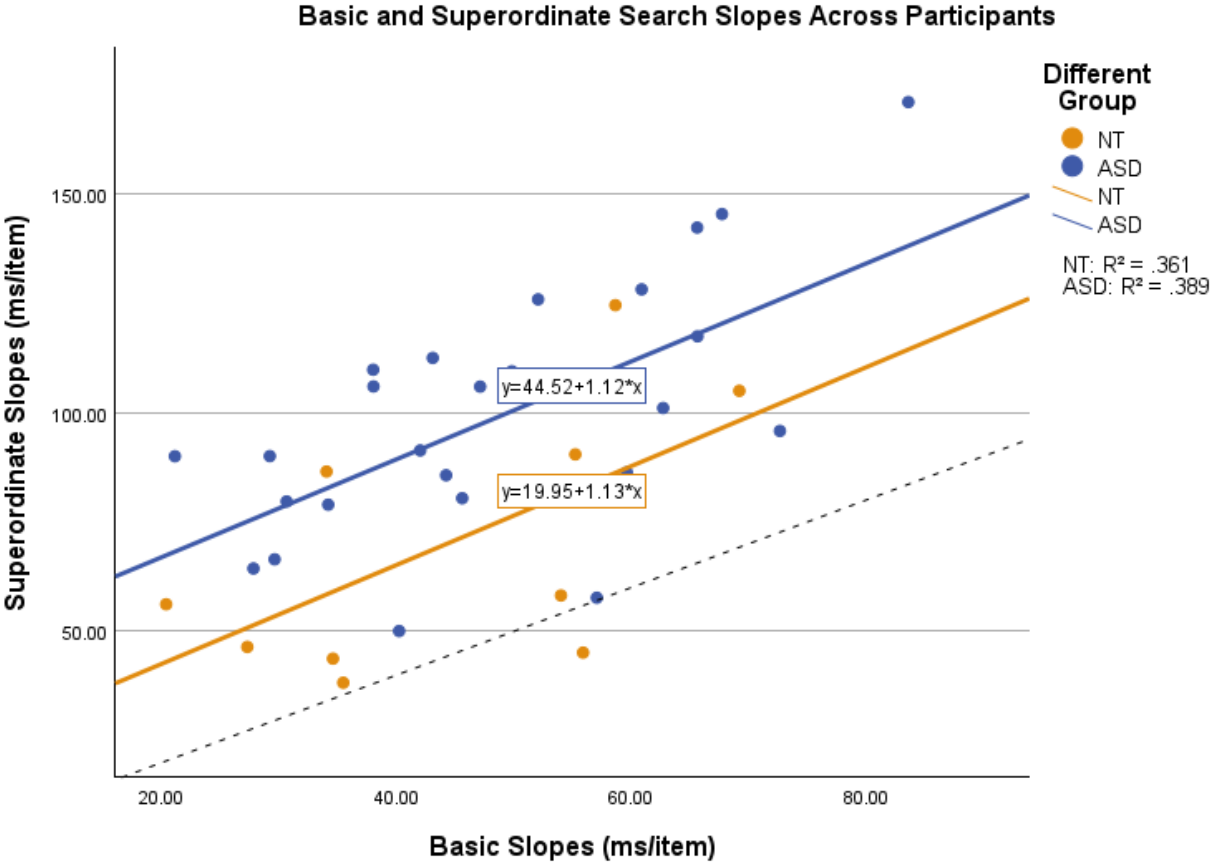
Basic- and superordinate-level search slopes in ASD (*n* = 26) and NT (*n* = 11) participants. Each point represents one participant, with basic-level slopes on the horizontal axis and superordinate-level slopes on the vertical axis, measured in milliseconds per item. The dashed black line marks equal slopes (*y* = *x*). Points above this line indicate steeper slopes for superordinate-level than basic-level search. R^2^ values describe the fitted relationships.

### Categorization

Reaction time in the categorization task was examined separately for the ASD and NT groups. Participants’ responses reflected the time required to select the member of a spoken category from a pair of images.

Item-level categorization times were compared between groups for vehicles, animals, food, kitchen tools and musical instruments (Figures 7 show examples). Across these categories, the displayed fitted lines had positive slopes: images associated with longer categorization times in the NT group also tended to have longer times in the ASD group. The points generally lay above the equality line, indicating longer categorization times in the ASD group. These comparisons describe the patterns shown in the figures.

**Figure 7.**
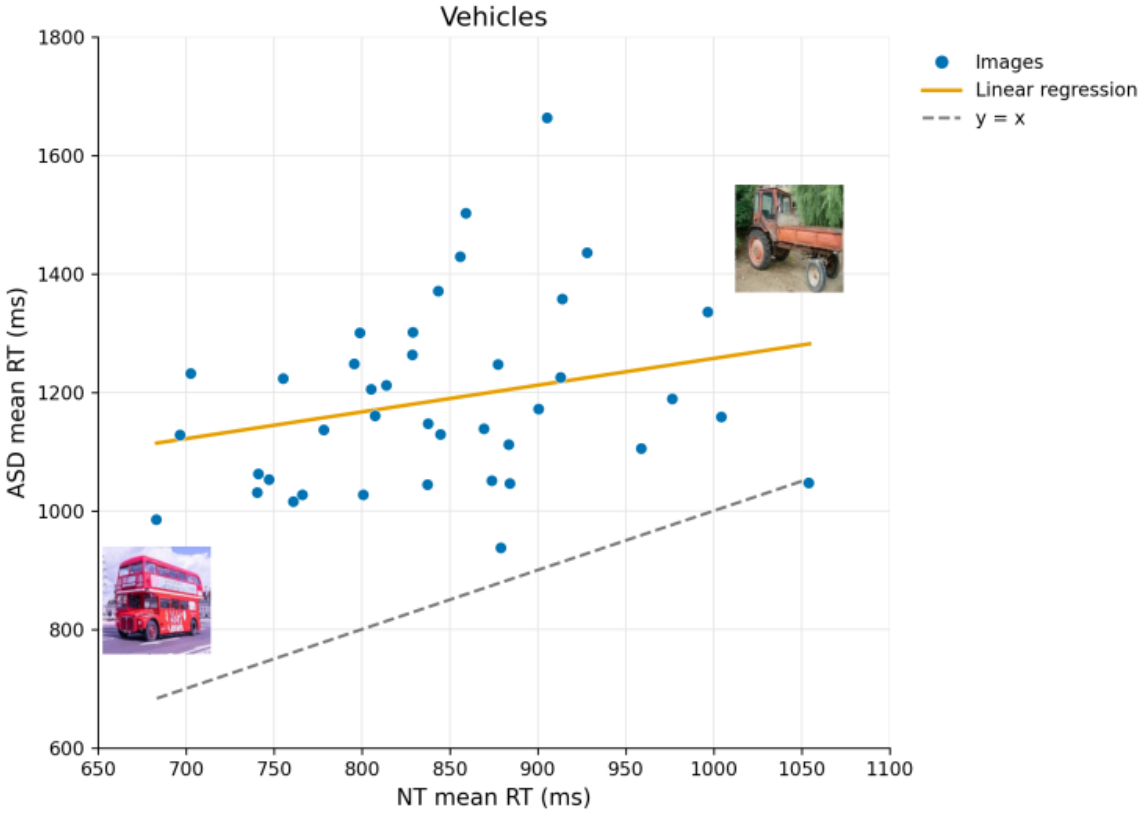
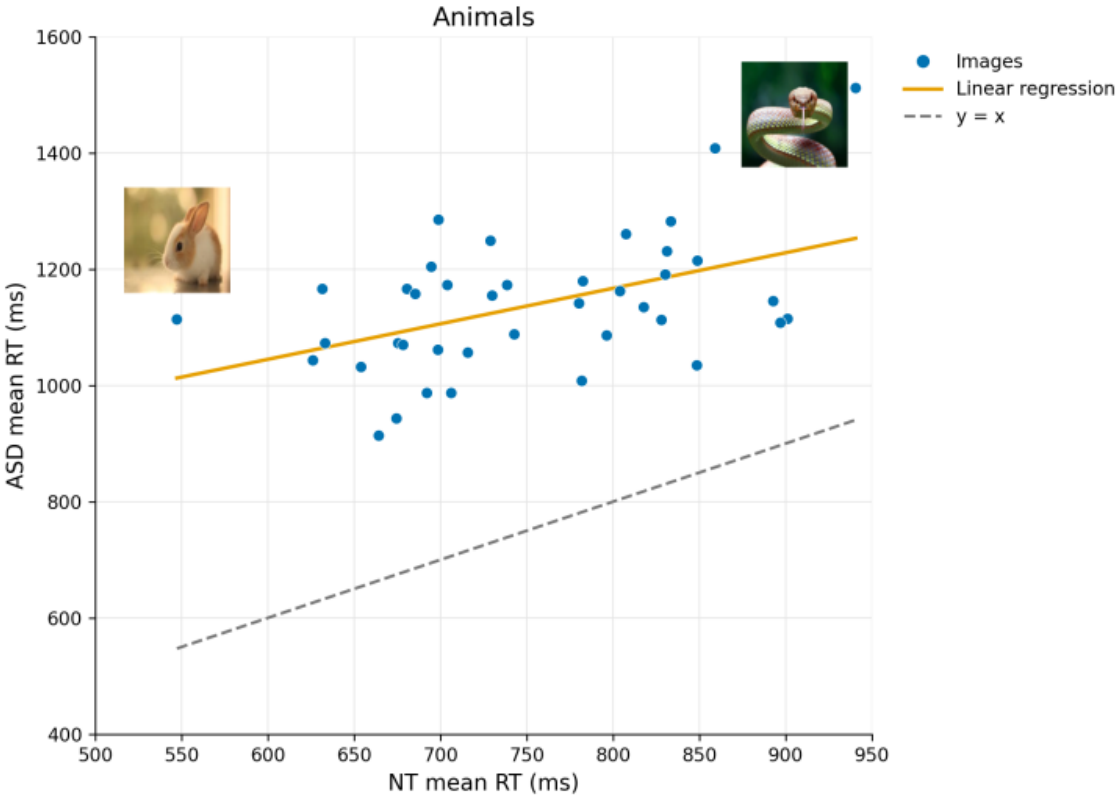
Item-level categorization times in the NT group (horizontal axis) and ASD group (vertical axis) for vehicles (top), animals (bottom). Each point represents an image. Solid lines show fitted relationships and dashed lines indicate equal RTs. Selected stimulus images illustrate examples.

### Relationship between visual search and categorization

To examine whether images found relatively quickly during visual search also elicited faster responses in the categorization task, reaction times for the same images were compared across tasks. Reaction times were normalized within each task to account for differences in their absolute values.

Figure 8 shows the relationship between normalized reaction times in the 8 × 8 visual-search condition and the categorization task across categories. Both groups showed a positive relationship, which was descriptively stronger in the ASD group (R^2^ = .143) than in the NT group (R^2^ = .026). Images found more quickly during visual search therefore tended to be categorized more quickly, although this correspondence was weak, particularly in the NT group. The difference in the strength of the associations between groups was not tested statistically.

**Figure 8.**
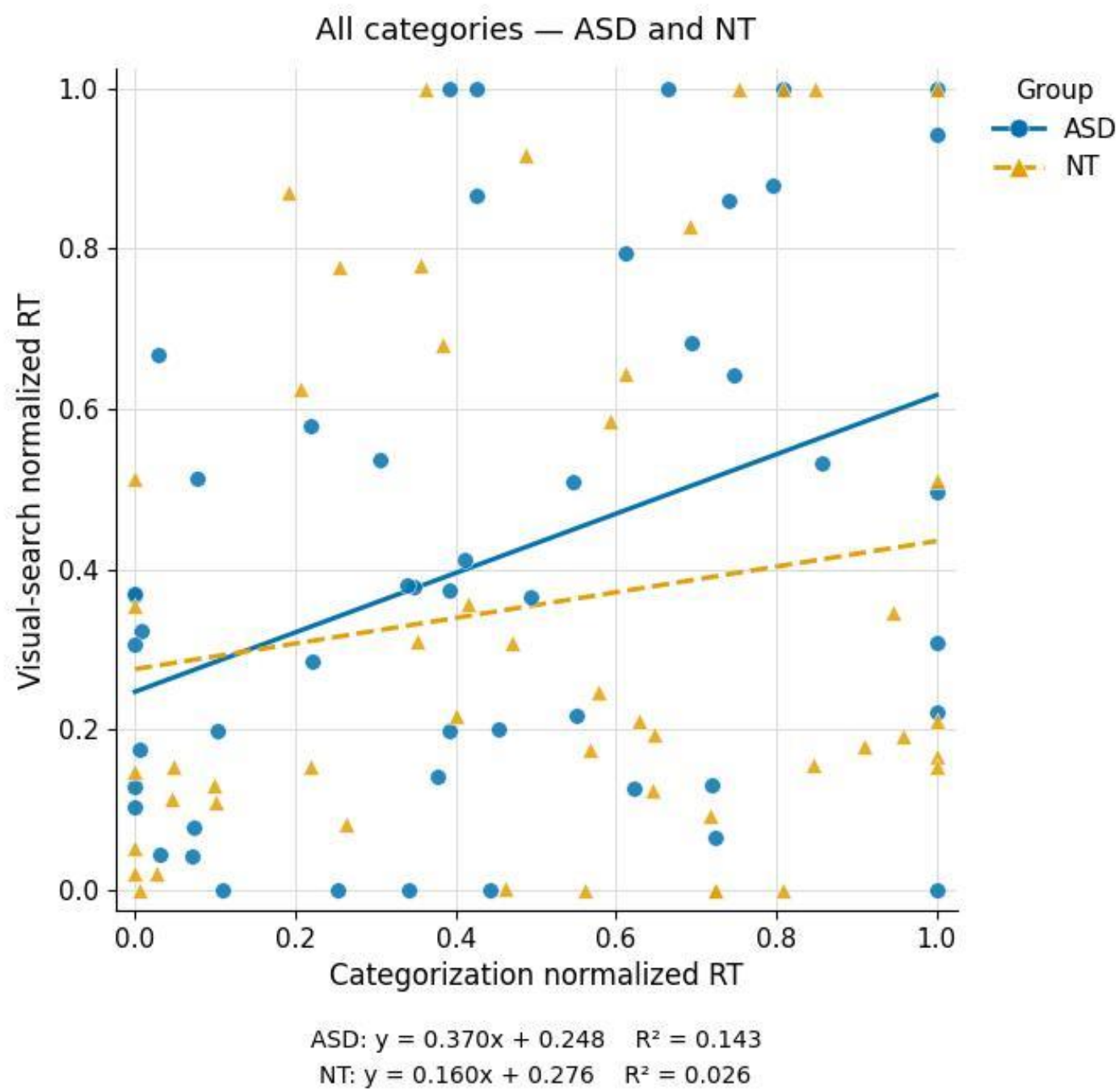
Relationship between normalized categorization reaction times (horizontal axis) and normalized visual-search reaction times in the 8 × 8 condition (vertical axis). Blue denotes ASD and orange denotes NT. Each point represents one image (ASD: R^2^ = .143; NT: R^2^ = .026)

## Discussion

The present study examined basic- and superordinate-level visual search in participants with autism and neurotypical participants, and the relationship between visual search and categorization. Both groups showed the same pattern: search slopes were shallowest for faces, followed by basic-level targets, and steepest for superordinate targets. However, moving from basic to superordinate search had a greater effect in the ASD group. Basic-level slopes were descriptively similar between groups, whereas the additional superordinate cost was approximately 50 ms/item in ASD and only 26 ms/item in NT. This suggests that searching for a broadly defined category placed greater demands on participants with autism.

A basic-level cue, such as “cat,” gives a more specific idea of what to look for than a superordinate cue such as “animal.” Schmidt and Zelinsky (2009) showed that more specific target cues provide more effective guidance during visual search. Maxfield and Zelinsky (2012) further demonstrated that category level affects search guidance and target verification differently. Related work by Hout and Goldinger (2015) showed that the precision of a target’s mental representation influences both attentional guidance and decisions about whether an object matches the target. Together, these findings suggest that a broad category cue may make it harder to direct attention toward possible targets and to decide whether an object belongs to the category. Either process could contribute to the larger superordinate search cost among participants with autism.

The relatively efficient face search in both groups is consistent with Abassi Abu Rukab et al. (2022) and it fits the face advantage described by Hershler and Hochstein (2005). It is also consistent with reverse hierarchy theory, which proposes that high-level category information can support rapid detection without requiring detailed visual scrutiny (Hochstein & Ahissar, 2002). However, the similar ordering of search difficulty does not establish that both groups used the same strategy. In both groups, participants whose search times increased more sharply with set size in basic-level search tended to show the same pattern in superordinate search. The relationship was similar in strength across groups, although it was statistically significant only in ASD. This suggests that individual differences in search efficiency were present at both levels, while the move to superordinate search added greater difficulty for the ASD group.

In the categorization task, images that took longer for the NT group also tended to take longer for the ASD group, with generally longer response times in ASD. Comparing the same images across tasks showed that some objects took longer both to find in the visual search task and to categorize in the 2AFC. This relationship appeared stronger in ASD, with the clearest category-level associations for kitchen tools and musical instruments.

The item-level comparisons showed that images that took longer for the NT group to categorize also tended to take longer for the ASD group, with generally longer response times in ASD. Comparing the same images across tasks suggested that objects that took longer to find also tended to take longer to categorize. This relationship appeared stronger in ASD, with the clearest category-level associations were found for kitchen tools and musical instruments. These findings suggest a possible link between visual search and categorization that warrants further investigation.

One possible explanation concerns how typical an object is of its category. Less typical objects may require closer examination before their category membership becomes clear, making them slower to find and categorize. Gastgeb et al. (2006) found that participants with autism took longer than matched controls to categorize atypical examples (see also Shulman, Yirmiya & Greenbaum, 1995). This makes typicality a relevant explanation to consider for the present findings. However, our participants selected a category member beside a distractor, and typicality was not measured independently. The slower responses could therefore also reflect familiarity, the visual features of the image, or difficulty distinguishing the target from its distractor.

Our findings raise a question for future research: do participants with autism need more time to find a possible target, to decide whether it belongs to the named category, or both? Combining eye tracking with reaction times could help us examine these stages more closely. We could also compare category names with picture cues to explore how the information provided before search affects performance. This would help us better understand how participants use category knowledge when searching for an object.

## Conclusions

The present study identifies category level as an important factor in understanding visual search among participants with autism. Basic-level search slopes were descriptively similar across ASD and NT groups, whereas searching for members of broader, superordinate categories produced a larger increase in slopes among participants with autism. This pattern suggests that group differences become more apparent when the task requires identifying targets that belong to the same category but differ in their visual features. The positive relationship between basic- and superordinate-level slopes also highlights individual differences in search efficiency within both groups.

The contribution of this study is to connect visual search with the category judgments required to identify a target. Examining performance across category levels helps clarify which search conditions are more challenging for participants with autism and shows why category level should be considered when interpreting their visual-search performance. It also raises a meaningful question for future research: how effectively can category knowledge guide attention when visually different objects belong to the same target category? Understanding this relationship could inform the design of learning materials and visual supports that make relevant category information easier to use.

## Acknowledgements

We thank all participants and their parents and teachers for facilitating this study. We thank the Ministry of Education Chief Scientist Office and the school principal for granting permission for this study. Thanks to Yuri Maximov for assistance with programming and data analysis.

This study was supported by a grant from the Israel Science Foundation (ISF) Corresponding Author: Shaul Hochstein

Address: Hebrew University, Jerusalem; ELSC & Life Sciences Institute, Safra Campus, Jerusalem Israel

